# An unsupervised deep learning framework encodes super-resolved image features to decode bacterial cell cycle

**DOI:** 10.1101/2024.03.13.584804

**Authors:** Juliette Griffié, Chen Zhang, Julien Denereaz, Thanh-An Pham, Gauthier Weissbart, Christian Sieben, Ambroise Lambert, Jan-Willem Veening, Suliana Manley

## Abstract

Super-resolution microscopy can resolve cellular features at the nanoscale. However, increased spatial resolution comes with increased phototoxicity, and reduced temporal resolution. As a result, studies that require the highest spatial resolutions often rely on static or fixed images, lacking dynamic information. This is particularly true of bacteria, whose lateral dimensions approach the scale of the diffraction limit. In this work, we present Enso, a method based on unsupervised machine learning to recover bacterial cell cycle and cell type information from static single molecule localization microscopy (SMLM) images, whilst retaining nanoscale spatial resolution. Enso uses single-cell images as input, and orders cells according to their spatial pattern progression, ultimately linked to the cell cycle. Our method requires no *a priori* knowledge or categories, and is validated on both simulated and user-annotated experimental data.

## Introduction

Super-resolution microscopy can produce images with spatial resolution below the diffraction barrier (∼200 nm), whilst retaining advantages of fluorescence microscopy: high target specificity on multiple biomolecules simultaneously (multi-color), and access to living samples (dynamics). This has proven especially powerful for bacterial cell biology, since an individual bacterium can be as small as 100-200 nm in linear dimensions. For example, single molecule localization microscopy (SMLM) in fixed cells exhibited clusters of proteins involved in chemotactic signaling whose size depended on the biochemical network state^1^. The role of major nucleoid-organizing proteins in global transcriptional silencing was dissected in living *Escherichia coli* cells^2^. Also via SMLM in living cells, the organization of the bacterial tubulin homologue, FtsZ, was observed as a band of semi-overlapping filaments in *Caulobacter crescentus*^3^. Yet, due to the limited number of protein copies and the large number of raw frames (hundreds to thousands) needed to reconstruct an image, time-lapse imaging with SMLM is rare: thus, these studies relied on single snapshots of individual cells.

Bacterial cell shape is largely determined by a rigid cell wall, which comprises cross-linked peptidoglycan (PG) strands. As cells double in volume during their cell cycle, the cell wall is remodeled to accommodate this growth. Curiously, different species have distinct patterns of remodeling, as revealed by the incorporation of a pulse of fluorescent d-amino acids (FDAAs)^4^. Models of cell shape dynamics based on such studies typically rely on length ordering of diffraction-limited snapshots, whereby the position of cells in cell cycle are presumed to vary directly with their length^5,6^. Structured illumination microscopy (SIM) can recover some temporal resolution, taking advantage of the lower light doses relative to SMLM, while doubling the resolution compared with widefield imaging. Taking this approach, a study of *Staphylococcus aureus* found elongated intermediate states and asymmetries in cell wall growth^7^. However, SIM does not offer the near molecular-scale resolution of SMLM, and no current methods exist to detect dynamic patterns in bacterial cell organization in a manner agnostic of cell shape or independent of user-defined shape parameters.

Considering other complex data types, an entirely different approach has been developed. Omics data sets such as from RNA or DNA sequencing consist of static, multi-dimensional representations -- similar to microscopy images. As a means to decipher them, unsupervised machine learning tools have been used to generate lower-dimensional, information enriched representations^8–10^. Remarkably, this approach has been successful at extracting reliable information about key cellular processes, including cancer subtype classification^11,12^. Furthermore, autoencoder frameworks can be extended to infer novel information such as drug treatment outcome or cell lineage from incomplete data sets^13,14^.

Here, we show that a similar approach can be applied to gain insight into SMLM image data sets. We developed a software tool predicated on a generative model architecture, which compresses image data into a commonly shared latent space. Our tool provides a general framework to investigate bacteria; because it is unsupervised it avoids user bias and does not require *a priori* knowledge or manual categorization. Our results demonstrate that dynamic processes such as cell shape progression linked to the bacterial cell cycle can be extracted from the interpretable latent space; thus, we call our method Enso (circle). We verified this approach on both simulated and experimental data sets of bacteria. We further demonstrate our approach can be applied to reveal patterns in data sets containing multiple cell types; distinguishing species whilst retaining this dynamic information.

## Results Workflow

Our goal is to develop a generalizable framework for extracting information on complex dynamic processes from images of fixed bacterial cells. To preclude reliance on *a priori* knowledge, we use generative models, a subgroup of unsupervised machine learning models which produce synthetic data sets. In the case of images, that translates into training the generative model on (experimental or simulated) input images which it “learns” to accurately replicate and output as synthetic, generated images. The model is considered trained when output and input images become indistinguishable (based on a one-to-one mapping loss estimate). Generative adversarial networks (GAN)^15^ and variational auto encoders (VAE)^16^ are examples of generative models that rely on deep neural networks.

Enso consists of a VAE whose architecture has been tailored to microscopy inputs. It is composed of two deep neural networks: the encoder and the decoder (**Fig. 1a**). The encoder is designed to compress the input data into a low-dimensional space, called the latent space. The decoder takes points in the latent space as inputs and decompresses them into images that should closely resemble the original set of images. Over multiple training epochs (cycles during which the training dataset is passed through the network), both encoder and decoder optimize their network weights to decrease the differences between the output and input images. We quantify the similarity between inputs and outputs by calculating the multi-scale structural similarity index measure (MS-SSIM)^17^, which is particularly suited to microscopy images because they contain information over multiple scales.

**Figure 1:**
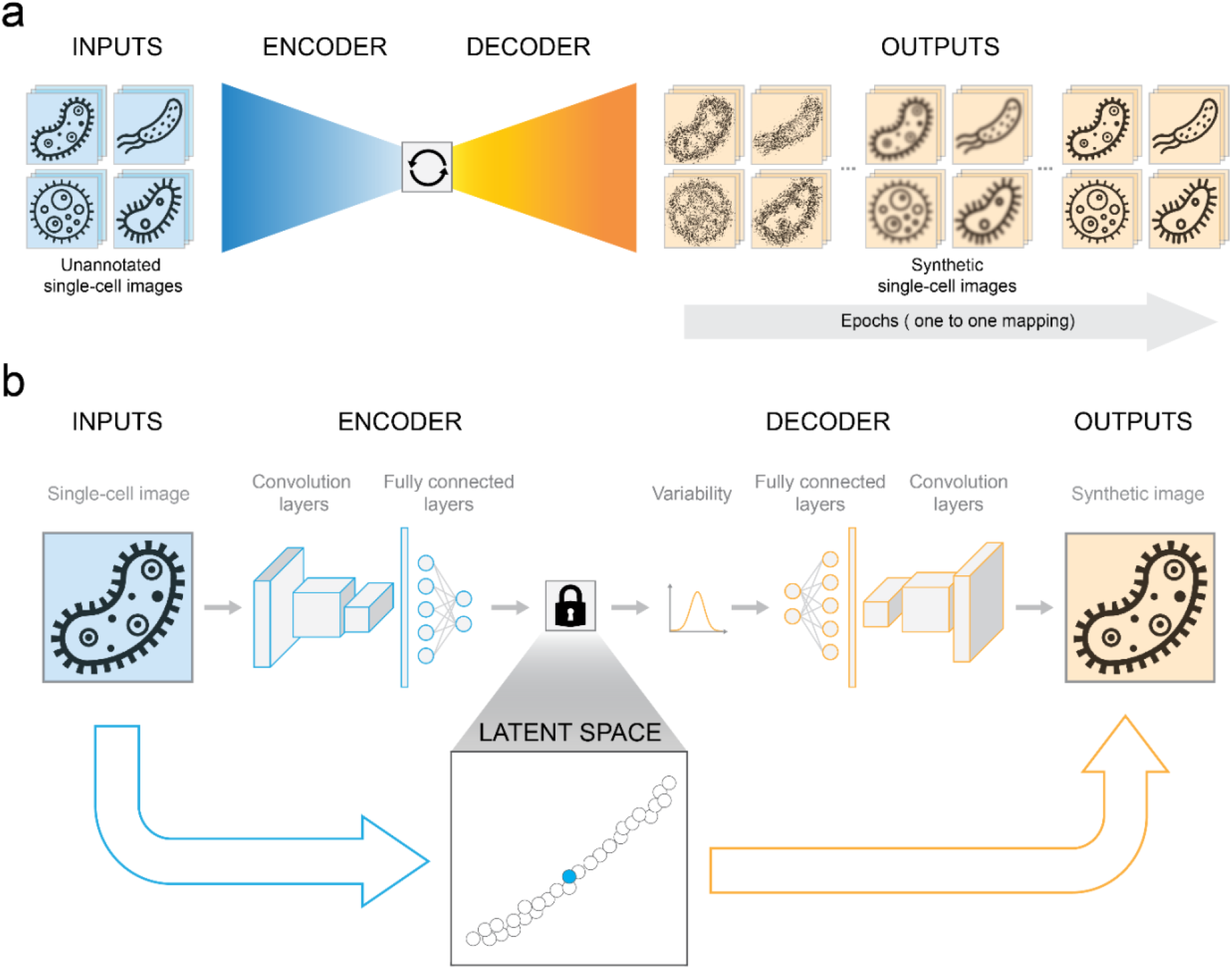
Enso workflow. (**a**) Training phase. The network “learns” to replicate input data sets (i.e., super resolved microscopy images). (**b**) Detailed layered architecture of Enso VAE. Unsupervised dimensionality reduction to a 2D latent space via convolution layers and fully connected layers.

The architecture of Enso (**Fig. 1b**) includes an encoder which uses convolutional layers to extract features from the images, which are captured in the latent space, then fully connected layers to perform dimensionality reduction. The decoder is designed to mirror the encoder. In contrast with the statistical technique of principal component analysis (PCA), which also performs unsupervised dimensionality reduction, Enso accommodates non-linearity and feature extraction through its convolutional layers. In the latent space, each image is converted into a coordinate for which the (integer) number of dimensions is pre-defined. The latent space representation hence consists of a cloud of points, with each point corresponding to an input image. The performance of a VAE in generating realistic synthetic data depends on how efficiently the latent space captures the essence of the data in a compressed fashion within its reduced dimensions. We tested several latent space dimensionalities: 1D, 2D, and 3D (**Fig. 2 and supplementary Fig. 1**). Because there was little difference in the MS-SSIM achieved by 2D and 3D, we focus on the 2D latent space as it is simpler to represent visually.

**Figure 2:**
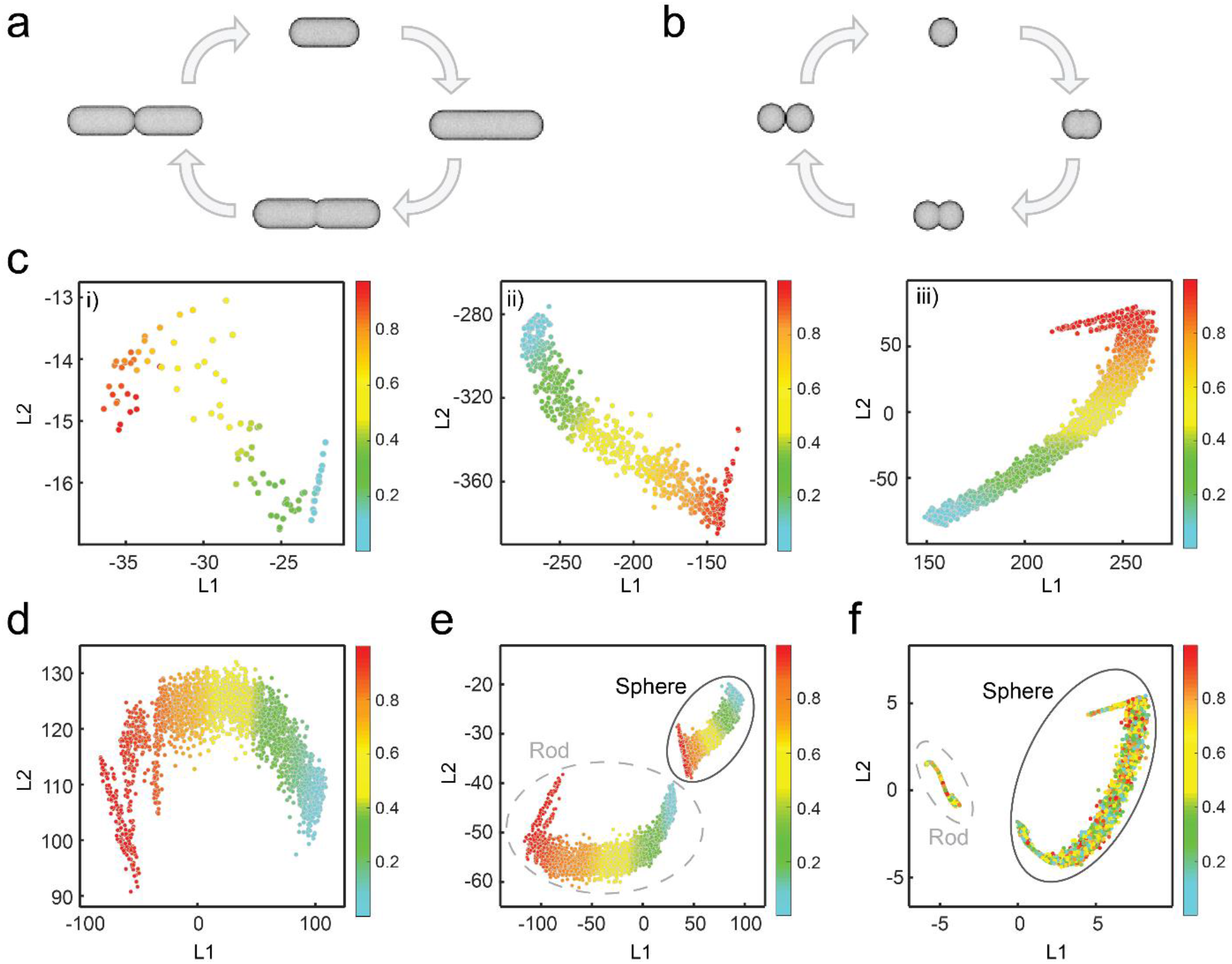
Enso performance on simulated data sets. Simulated cell cycle in (**a**) rod shape model (**b**) spherical shape model. (**c**) Rod shape: latent space representation after training 50 epochs for 100 (i), 750 (ii), and 3000 (iii) single cells as input. (**d**) Spherical shape: latent space representation after training 50 epochs for 3000 input images. (**e**) Enso latent space representation when training with a mixed data set composed of rod- and spherical-shaped bacteria images. (**f**) Principal component representation when applied to a mixed data set composed of rod shape and spherical bacteria images.

### Simulated data sets

Since we anticipated that microscopy images would contain bacterial cells at different stages of the cell cycle, we built a simulated model of cell growth to aid in interpretation. The simulations consist of images of 144 x 64 pixels (for equivalence with experimental images, pixel size = 30 nm). Each image contains a single cell, and similar to images acquired with a physical microscope, its stained cell wall is projected into 2D. In our model, newly born cells just after division at time t = 0 were assigned a length, then allowed to grow over a period of time t_d_, before dividing. We produced bacillus (rod)-shaped bacteria whose lengths increase from 1000 to 2000 nm over the cell cycle, and whose widths vary from the cell diameter of 700 nm to fully constricted (zero) just before division (**Fig. 2a**). By randomly selecting simulated images t ϵ [0, t_d_], we generated a population of static images equivalent to fixed cells at different stages of the cell cycle. We retained the information on time to use it as the ground truth for cell cycle dynamics throughout our analysis of simulated data (**Fig. 2**).

The workflow uses simulated images as input, and produces a corresponding synthetic image from the VAE after training, where each input image resides as a point in the resulting latent space. We populated a 2D latent space with increasing numbers of input single-cell images (ranging from 100 to 3000) (**Fig. 2c**). In the latent space representation, we color-coded each point by the ground truth i.e., the *t* value representing its cell-cycle stage. We observed that within the latent space, cell images appear ordered by *t* automatically. By compressing each data set within the reduced dimensionality latent space, we recovered the dynamic cell cycle order from individual snapshots. Crucially, this was done in an unsupervised manner: no user inputs were required for this task, unlike classical methods of pseudo-time ordering for bacteria which sort cells according to length. Similarly, cell-cycle inference is retained for a 1D or a 3D latent space (**Supplementary Fig. 1**).

Furthermore, our results highlight that even with a relatively small training set (i.e., 100 images), cells ordered accurately within the latent space (**Fig. 2c**). Enso has minimal computational requirements, in contrast to other deep neural network-based methods that require access to GPUs and a considerable number of images as inputs, often a deterrent to using such methods. Since our method can be implemented using CPU, minimal technical skills are required to set it up, making it more accessible to non-expert users including biology-focused labs. All of the results reported here were generated on a conventional PC with computation time ranging from a few minutes to a few hours.

Using our simulator, we could examine the influence of different noise sources on Enso’s capacity to perform ordering along the cell cycle. We varied staining density and signal to noise ratios, common issues with SMLM and other types of super-resolution images. We found that our method produces an ordering that is highly robust to these factors (**Supplementary Fig. 2, 3**).

Bacteria come in several shape archetypes, so we also applied Enso to study coccus-shaped (spherical) cells. We produced images of cocci whose length varied from 700 to 1400 nm over the cell cycle, and whose widths vary from 700 nm to fully constricted (zero) just before division (**Fig. 2b**). We then used the same VAE architecture, and found that cocci also ordered according to their time in cell cycle within the latent space (**Fig. 2d**). Since environmental samples or microbiome-like studies contain cells of mixed type co-existing together, we wondered how Enso would perform on more complex mixtures of input images. By training the network on an input data set including both cell types (i.e., bacilli and cocci), we observed that the latent space representation retained the dynamic cell-cycle information; moreover, the two cell shapes clustered separately into two distinct point clouds (**Fig. 2e**). When analyzed with a PCA approach, cell types could be distinguished; however, all dynamic information was obscured (**Fig. 2f**). We suspect that this is due to PCA’s insensitivity to nonlinearities in the data.

### Experimental data sets

We then tested Enso’s performance using real biological data sets. Crescent-shaped *C. crescentus* and ovoid-shaped *Streptococcus pneumoniae* cells were cultured, labeled, plated on agarose pads and imaged using a large field of view SMLM setup^18^. Individual cells were segmented from each field of view, aligned and converted into images of 64 x 192 pixels (pixel size = 30 nm). We then trained Enso on two data sets separately: *C. crescentus* (**Fig. 3a**. i-ii, N = 2396 cells), and *S. pneumoniae* (**Fig. 3b**. i-ii, N = 2189 cells). Cells within each latent space appear as a cloud of points (grey), with each point representing a single cell (**Fig. 3a, b**. iii). The model outputs synthetic cell data which closely resembles the input data, as desired (**Fig. 3a, b**. iv).

**Figure 3:**
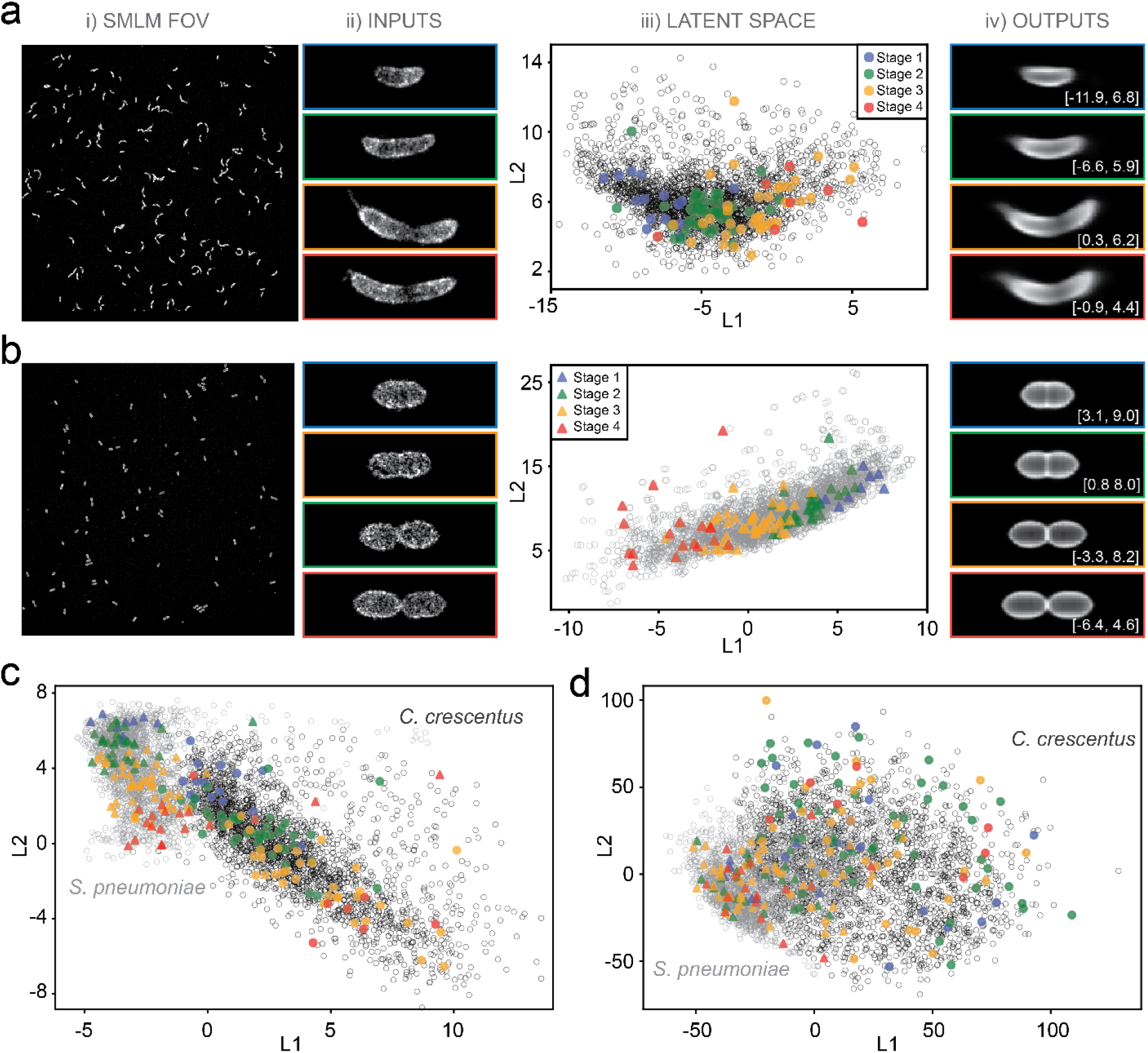
Validation on Experimental data sets. (**a**) Results on mixed population of C. crescentus cells, with an example field of view (FOV) (i), segmented cell images used for training (ii), the latent space with all cells in grey and manually annotated cells in color according to cell cycle stage (iii), and network outputs with their position in the 2D latent space [L1, L2] (iv). (**b**) Results on mixed population of S. pneumoniae cells. (**c**) Results on mixed cell types data sets using Enso (triangles, S. pneumoniae; circles, C. crescentus). (**d**) Results on mixed cell types data sets using PCA.

To be able to interpret patterns within the latent space, we randomly selected 5% of the images from each data set and manually labeled them to approximate the ground truth cell cycle stage (**Supplementary Fig. 4**). When plotted within the latent space, we observed that, as for simulated data sets, experimental data sets produced an ordered representation along the cell cycle (**Fig. 3a, b**. iii). *C. crescentus* has a complex cell cycle since it undergoes asymmetric cell division and differentiation from a flagellated swarmer cell into a stalked cell. This may explain the noisier ordering and the broader distribution within the latent space compared to *S. pneumoniae*, a cell type which presents less variability. This dynamic information is also retained in the case of a 1D or 3D latent space (**Supplementary Fig. 5**).

Next, we trained Enso with a mixed data set of *C. crescentus* and *S. pneumoniae* cells (**Fig. 3c**, N = 4585 cells). When both cell types are used as input, each cell type appears as a well-separated cluster whilst continuing to order along its own cell cycle (**Fig. 3c**). In contrast, PCA taking as inputs the same data sets results in strong overlap between species, and lacks dynamic information (**Fig. 3d**). Using cluster analysis for spatial point patterns (SPP)^19^, we moreover assessed that Enso accurately attributes 94% of provided cells to the correct species (72% with PCA), 1% attributed to the wrong species (27% for PCA) and 5% non-attributed data points (1% for PCA). Overall, these results suggest that Enso is capable of extracting dynamic features from static experimental images containing real noise, as well as cell type identification.

*C. crescentus* has been developed as an experimental model system for studying the cell cycle, due to its amenability to synchronization. Synchronization is performed by density centrifugation, which isolates swarmer cells at a similar early cell cycle stage (G0). Following this approximation of t = 0, cells resume cell-cycle progression and can be studied and imaged over time. As such, synchrony is routinely used to approximate the ground truth when it comes to the cell cycle (i.e., it is assumed that post-synchrony the cells are all at the same cell stage). A common practice in the data analysis of synchronized cells is population averaging, whereby protein organization, genetic profiles, or shape-associated features are averaged across many cells at equivalent times post-synchrony. However, using Enso we find that within a synchronized population, for over a third of the cell cycle, two populations of cells are present, including cells which have already divided (**Extended Data Fig. 6**).

Growth in rich media leads to faster cell doubling and a shorter cell cycle whilst growth in poor media leads to slower cell doubling and a longer cell cycle. Unlike other model systems such as *E. coli* and *Pseudomonas aeruginosa* which change their size and shape depending on nutrient conditions, *C. crescentus* appears to maintain its overall morphology. Studies suggest that the cell cycle rate changes, but control and homeostasis are maintained by similar mechanisms, giving rise to a scaling relation with respect to the cell cycle duration^20^. To examine this finding, we imaged *C. crescentus* cells that were cultured in three different cell growth conditions: poor (M2G), standard (M2G), and rich (PYE) media. A single Enso model was trained using cells from the three different growth conditions as inputs. As expected, we find that there is no significant difference between the three growth conditions (**Extended Data Fig. 7**).

## Discussion

Enso enables unsupervised analysis of microscopy images, without any *a priori* knowledge. It is furthermore robust to noise (both random and biological) and amenable for use with a low number of training images. For these reasons it is suited to microscopy techniques for which throughput is otherwise a limiting factor to access deep neural network analyses. Its latent space representations encode biologically relevant information such as cell type and cell cycle dynamics, otherwise difficult to access from fixed cell images without a model and parametrized analysis. We show that it outperforms existing unsupervised, but linear approaches to image classification (i.e., PCA).

We anticipate that frameworks such as Enso presented here can offer dynamic insights in the case of extremophiles, anaerobes, or highly pathogenic cells for which live-cell imaging is particularly challenging. Such bacteria can have long cell cycle durations (e.g. a full day for *Mycobacterium tuberculosis*), resulting in low data collection throughput. Similarly, it enables characterization of cell types within a mixed population, and could prove useful when investigating biofilms present in natural and clinical settings. Since we have deployed Enso to analyze cells whose shapes are stereotyped, interpreting the latent space representation of other, more variable systems such as mammalian cells would require additional investigation. However, for yeast and bacterial cells with stereotyped shapes but more complex life cycles including differentiation, Enso may enable reconstructing the underlying trajectories.

In summary, Enso provides a data-driven framework for cell cycle inference, paving the way for the investigation of complex dynamic biological processes.

## Methods

### Bacterial cell culture and labeling

#### Caulobacter crescentus

Liquid *C. crescentus* CB15N cultures were grown overnight at 28 °C in 3 ml of M2G medium (M2 salts supplemented with 0.2% glucose)^21^ under mechanical agitation (200 r.p.m.). Liquid cultures were re-inoculated into fresh M2G medium to grow cells until log-phase (OD600 = 0.2–0.4). Cells were collected and fixed by 1 mL of 4% PFA in PBS (Alfa Aesar #Y13G502) at room temperature for 15 min, then washed with PBS and permeabilized by 1 mL of 0.2% TritonX-100 (ACROS Organics #A0413137) at 4 °C for 30 min. After two washes, cells were incubated by 1 mL of PBS containing 100 μg/ml WGA-conjugated Alexa Fluor-647 (Invitrogen #W32466) for 30 min, in darkness, to stain cell walls. Labeled cells were washed once with PBS, post-fixed by 1 mL of 4% PFA for 15 min as before, and resuspended in PBS. When using different culture mediums (PYE^21^ or M2X, M2 salts supplemented with 0.2% xylose), the labeling procedures remain the same as described above.

Synchronization of *C. crescentus* cells followed a previously described protocol^22^. In brief, mid log-phase cell cultures (OD_600_ ∼0.5) were centrifugated for 15 min, 7000 g at 4 °C. The supernatant and loosely pelleted cells were discarded. Pellets were resuspended and washed twice with cold M2 salt buffer.

Resuspension was mixed with cold Percoll (Sigma #P1644) with 1:1 volume ratio, and then centrifugated for 20 min, 15000 g at 4 °C. The bottom swarmer band (two bands should be observed) was collected and washed twice with M2, 15000 g for 3 min at 4 °C. The resultant pellets were resuspended into fresh M2G medium at OD_600_ ∼0.2. Cells grew post-synchrony were collected every 30 min for the labeling.

#### Streptococcus pneumoniae

Liquid *S. pneumoniae* D39V cultures were grown overnight at 37 °C in 3 ml of C+Y medium^23^ under static conditions. Liquid cultures were re-inoculated into 10 ml fresh C+Y medium to grow cells until early log-phase (OD_600_ ∼0.1). Cell pellets were collected by centrifugation and resuspended into 3 ml fresh C+Y medium containing 100 µM sCy_5_DA dye for cell wall labeling^24^. After ∼2 h (3-4 cell cycle durations), cells were collected, washed three times with 1 mL of PBS, and fixed by 1 mL of 4% PFA in PBS for 15 min at room temperature. Cells were then resuspended in 1 mL of PBS before imaging. For pulse labeling, re-inoculated *S. pneumoniae* cultures were grown until log-phase (OD_600_ ∼0.3), and then resuspended into 3 ml fresh C+Y medium containing 100 µM sCy_5_DA dye. After ∼15 min, cells were collected, washed, and fixed as the same as mentioned above. To break cell chains/clusters, bead-beating (without adding beads) was performed for post-fixed cells for 1 cycle of 30 seconds.

### Sample preparation

The SMLM imaging of bacteria was performed on an agarose pad made by 1x dSTORM buffer, as previously described^25^. In brief, the labeled bacteria were resuspended into 1x dSTORM buffer and spotted on the agarose pad for STORM imaging. The 1x dSTORM buffer was made by 20 mM sodium sulfite and 10 μM mercaptoethilamine (MEA) in PBS^26^. To make the agarose pad, 1 mL 2x dSTORM buffer (twice concentration of sodium sulfite and MEA) and 1 mL 3% agarose solution in ddH_2_O were mixed gently, and ∼400 μL of the mixture was added into a silicon gasket (Grace Bio-Labs #3110) placed on a rectangular glass slide (Epredia #0781). Another glass slide was then used to cover and seal the gasket from the top. The slides/agarose pad were kept in fridge (4 °C) for ∼20 min to let agarose fully solidified. The top slide was then removed, the gasket and agarose pad were carefully transferred onto a #1.5 round coverslip (25 mm in diameter, VVR #6310172) that fits the size of a custom-made holder. Finally, 1-2 μL drops of the cell suspension in 1x dSTORM buffer was spotted onto the pad. After a full absorption, the agarose pad was sealed with a plasma-cleaned #1.5 round coverslip (same size as the bottom one) from the top.

### Image acquisition

A custom built STORM microscope was used in this study^18^. For the image acquisition, a wide-field (WF) image was captured (200 ms exposure, 6.8 W/cm2) using the 642 nm laser line, followed by STORM imaging acquisition where 20000-30000 frames were taken (10 ms exposure, 3.4 kW/cm2). The 405 nm activation laser power was manually turned on from the minimum power intensity, and gradually increased when less photoblinking was shown. After acquisition, the raw STORM imaging stacks were processed by the ThunderSTORM^27^ plugin in Fiji^28^. The multi-emitter fitting analysis was enabled with a maximum of three molecules per fitting region. Drift correction was achieved by the cross-correlation tool. Duplicates that converged to the same localization uncertainty were removed. Images were visualized by averaged shifted histograms with a pixel size of 21.2 nm.

### Formatting

SMLM acquired image stacks were processed using ThunderSTORM. The resulting SPPs composed of fluorophores localizations were translated into pixelated FOVs (pixel size = 33 nm). Each individual cell was segmented from the FOVs using an intensity-based mask.

An intensity threshold is first applied at the pixel level over the original FOV. By this means, we binarize the FOV, generating a mask composed of pixels with values set at either 1, if above threshold, or 0 otherwise. The mask is then further processed by filling up existing gaps (i.e., a null pixel in contact with more than one pixel whose value is one, will be set to one as well). The resulting mask is composed of a number of objects (i.e., connected pixels of value 1) and null pixels. We use these detected objects to segment cells in the original FOV. Each segmented cell is then centered and flattened on a ROI of 64 x 192 pixels. The pixels values are then normalized over the ROI. These ROI are used as input for training and analysis with Enso.

### Manual labelling

For experimental datasets, since we could not directly access cell cycle stage ground truth, we manually labelled 5% of the data (2 annotators, overall results averaged at a single cell level). For *C. crescentus* (similarly for *S. pneumoniae*) we relied on 4 identified cell stages: 1-new cell, 2-elongation, 3-constriction, 4-dividing.

## Acknowledgements

We thank D. Sage for technical support. This work was supported by the Marie Skłodowska-Curie Fellowships (890169 BALTIC, to J.G.) and the European Union’s H2020 program under the European Research Council (ERC; CoG 819823 Piko, to S.M. and C.Z.).

## Contributions

J.G., S.M., A.L., and C.S. conceived and designed the project. J.G., S.M. and J.W.V. supervised the project. J.G and T.A.P. designed and implemented the VAE software package. J.G. designed and implemented the simulation software package. C.Z, J.G., J.D., A.L. and C.S. designed and optimized the experimental protocols. C.Z. collected the *C. crescentus* and *S. pneumoniae* data sets with assistance from J.D. and J.W.V. J.G., S.M. and C.Z. wrote the manuscript with the feedback of all coauthors.

---

**Supplementary Figure 1:**
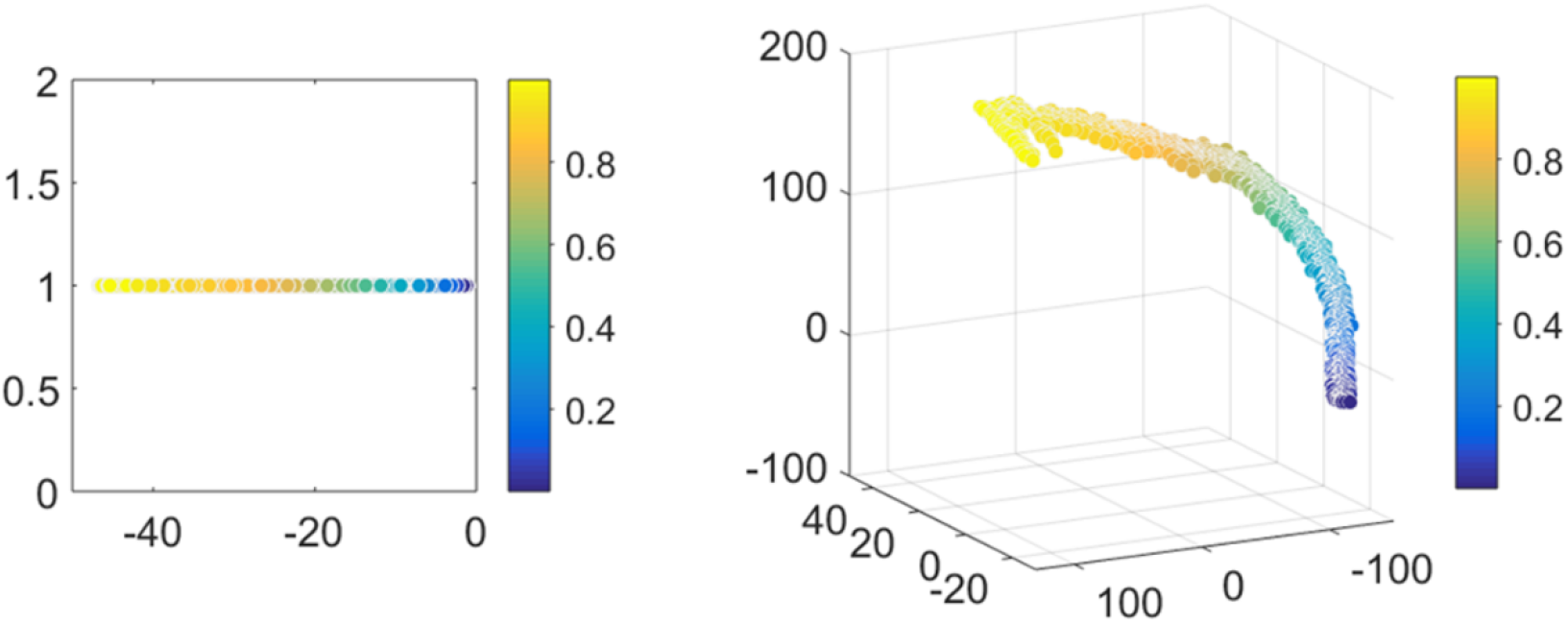
Enso ordering robustness to latent space dimensions for rod-shaped simulated bacteria.1D latent space (left) and 3D latent space (right).

**Supplementary Figure 2:**
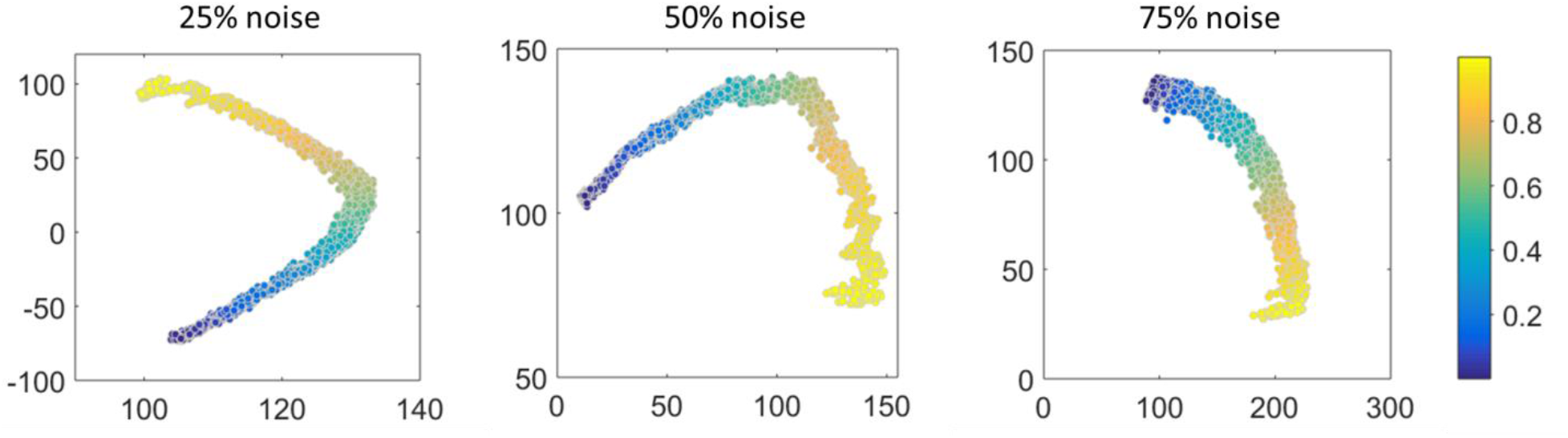
Enso ordering robustness overlaid random noise for rod-shaped simulated bacteria.

**Supplementary Figure 3:**
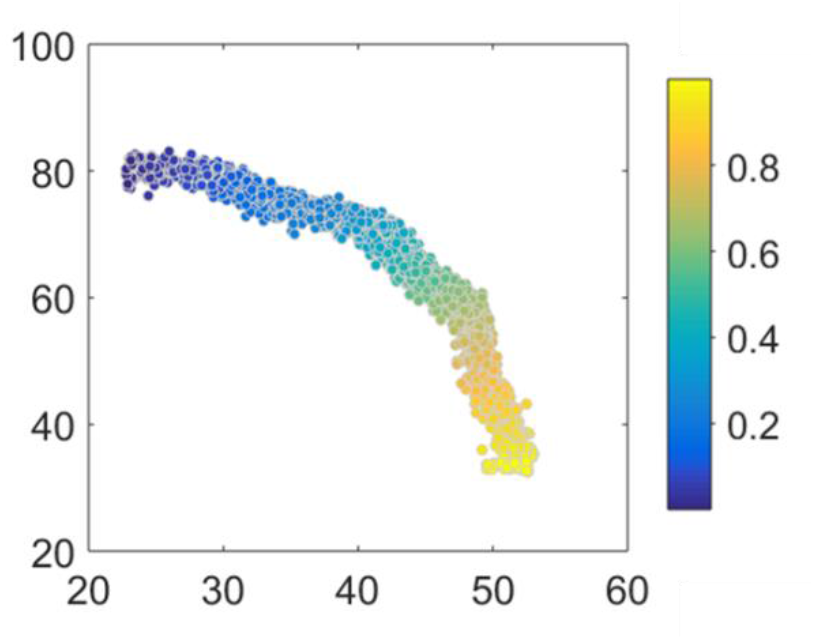
Ordering robustness to varying labelling density (over an order of magnitude). Rod shape simulated bacteria.

**Supplementary Figure 4:**
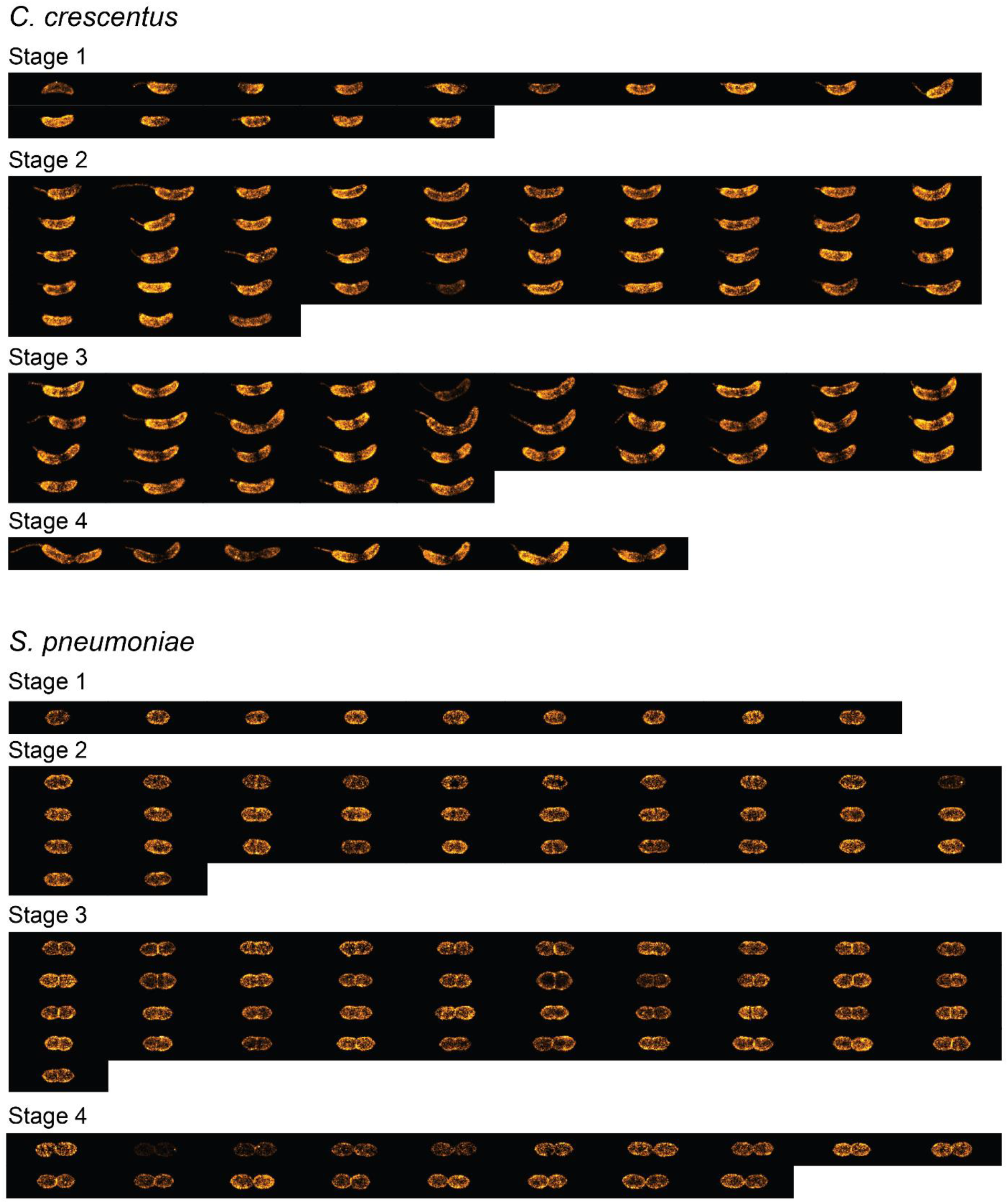
Manual labeling of 4 different stages of mixed *C. crescentus* and *S. pneumoniae* cells. N=100 in total for both species.

**Supplementary Figure 5:**
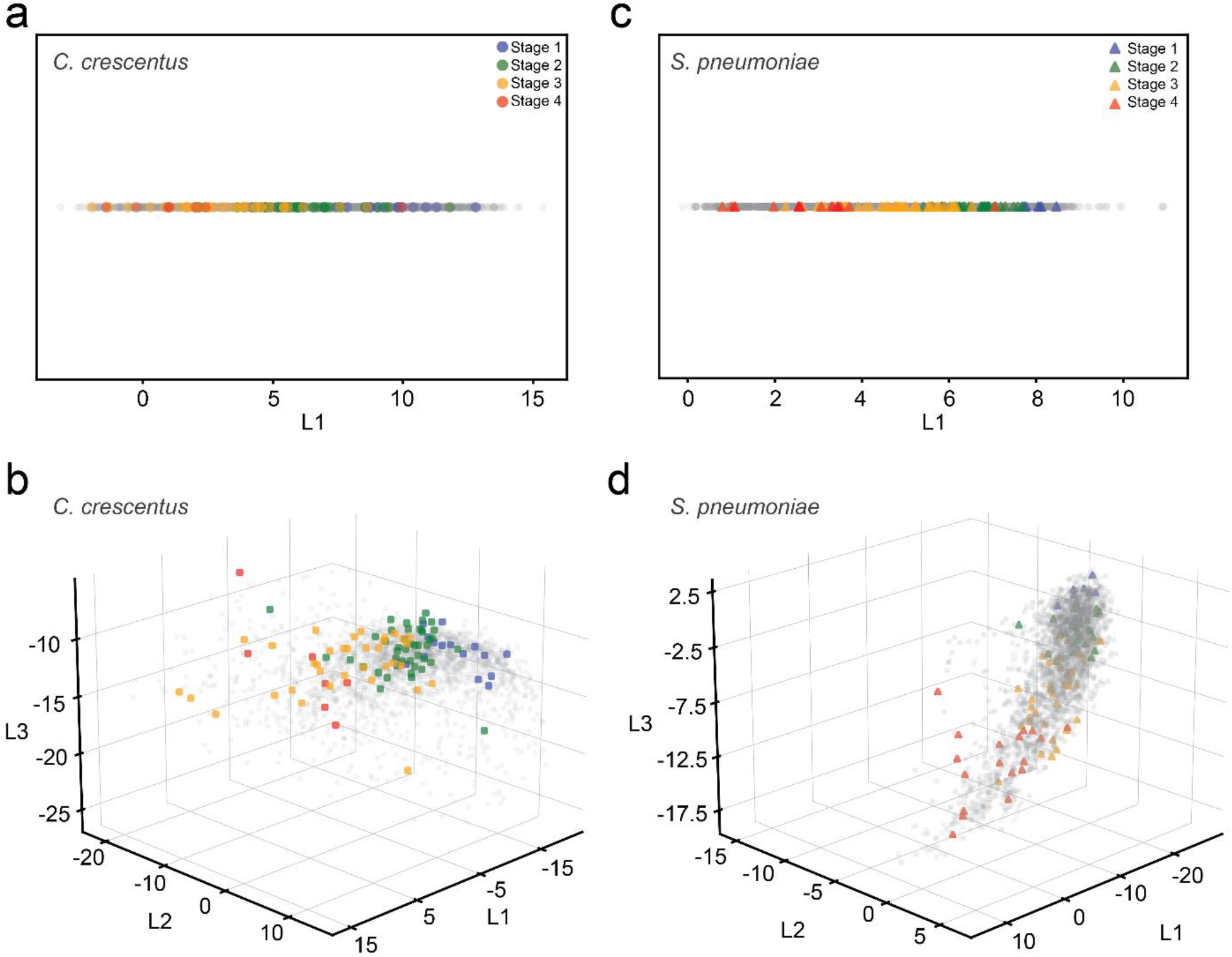
Enso ordering performance of *C. crescentus* and *S. pneumoniae* cells using 1D (upper) or 3D (below) dimension of the latent space representation.

**Supplementary Figure 6:**
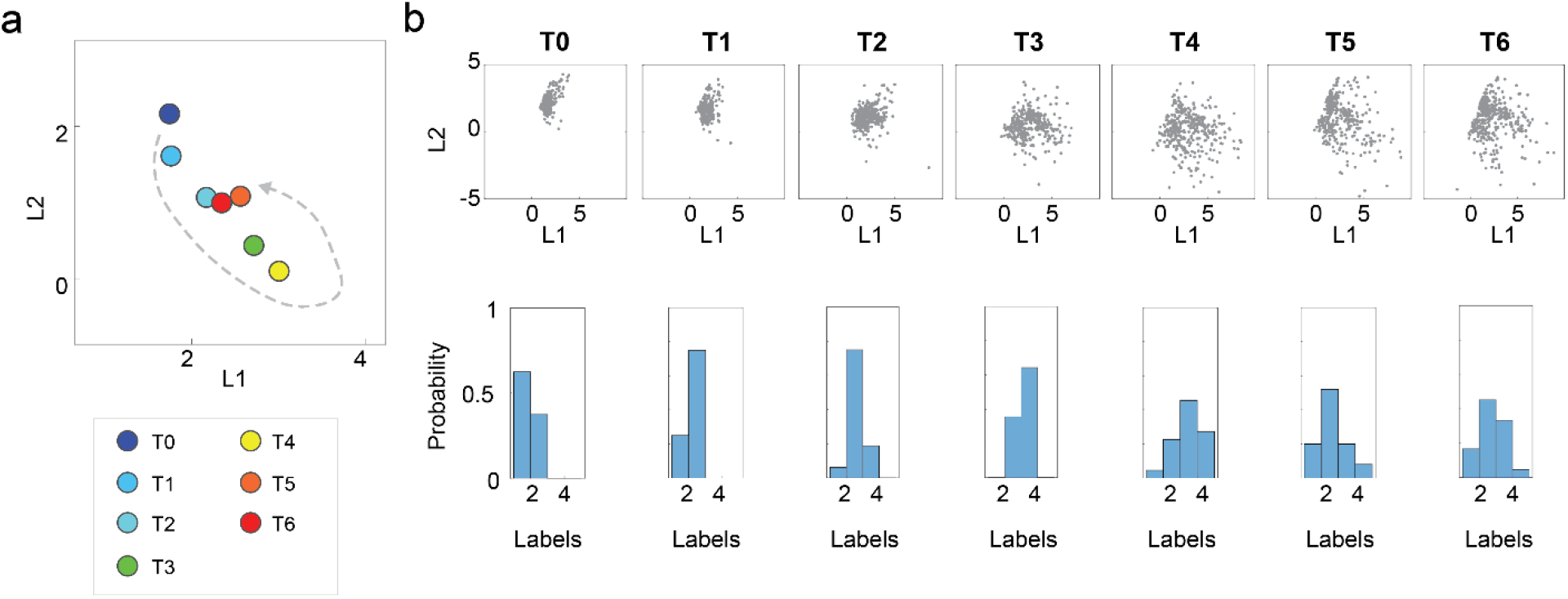
*C. crescentus cell* synchronization analysis. (**a**) Each point representation the mean coordinates in the latent space representation of the cloud of points associated to cells fixed at the same time point after synchronization. (b) Resulting cloud of points and histogram of cell stage (manual labelling of 4 cell cycle stages) obtained for cells at each time point following synchronization.

**Supplementary Figure 7:**
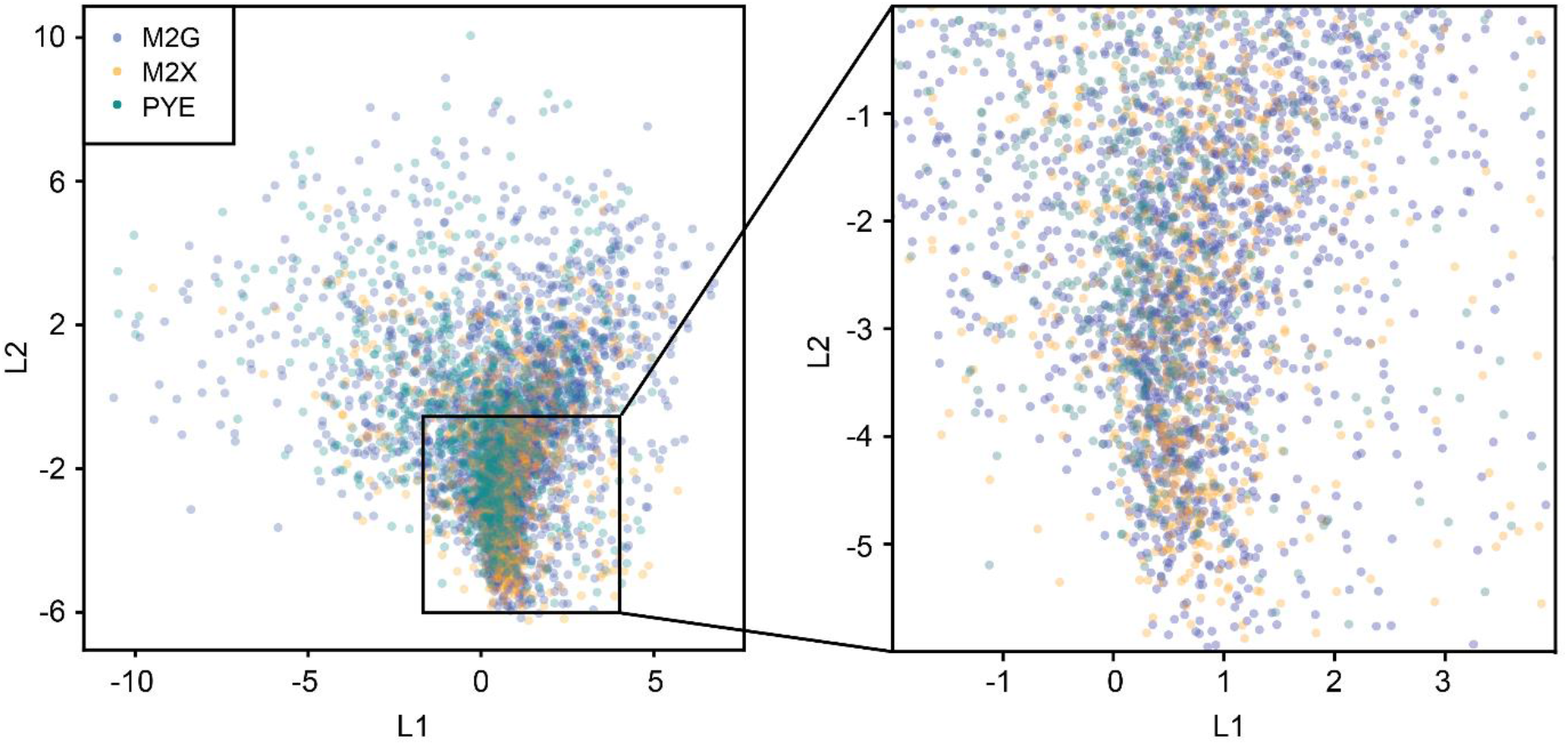
Latent space obtained for a model trained with a data sets composed *C. crescentus* segmented cells images cultured in 3 different cell growth conditions: M2G (poor), M2X (standard), PYE (rich).

